# Talin Rod Domain Containing Protein 1 (TLNRD1) is a novel actin-bundling protein which promotes filopodia formation

**DOI:** 10.1101/2020.05.19.103606

**Authors:** Alana R Cowell, Guillaume Jacquemet, Abhimanyu K Singh, York-Christoph Ammon, David G Brown, Anna Akhmanova, Johanna Ivaska, Benjamin T. Goult

## Abstract

Talin is a mechanosensitive adapter protein which couples integrins to the cytoskeleton and regulates integrin-mediated adhesion. Talin rod domain-containing protein-1 (TLNRD1) shares 22% homology with the R7R8 domains of talin, and is highly conserved throughout vertebrate evolution, however little is known about its function. Here we show that TLNRD1 is an α-helical protein which shares the same atypical topology as talin R7R8, but forms a novel antiparallel dimer arrangement. Actin co-sedimentation assays and electron microscopy reveal TLNRD1 is an actin-bundling protein that forms tight actin bundles. In addition, TLNRD1 binds to the same LD-motif containing proteins, RIAM and KANK, as talin, and thus may act in competition with talin. Filopodia are cell protrusions supported by tightly bundled actin filaments and TLNRD1 localises to filopodia tips, increases filopodia number and promotes cell migration in 2D. Together our results suggest that TLNRD1 has similar functionality to talin R7R8, serving as a nexus between the actin and microtubule cytoskeletons independent of adhesion complexes.

## Introduction

Talin rod domain-containing protein 1 (TLNRD1) is an evolutionarily conserved yet little studied protein that shares high sequence homology with the central region of the cytoskeletal protein talin. TLNRD1 was originally named Mesoderm Development Candidate 1 (MESDC1) because the gene, located on human chromosome 15, mapped to the *mesd* locus essential for mesoderm development (Holdener et al., 1994; Wines et al., 2001). This region contains two genes, *Mesdc1* and *Mesdc2*, and further analysis showed that the *Mesdc2* gene product, now renamed to MESD, was essential for mesoderm differentiation and embryonic polarity (Hsieh et al., 2003; Holdener-Kenny et al., 1992). In contrast, the *Mesdc1* gene was unable to rescue the lethal phenotype produced by deletion of the *mesd* gene region, suggesting the name was misleading. Thus, MESDC1 has been largely ignored, and it was renamed <u>T</u>a<u>l</u>i<u>n</u> <u>R</u>od <u>D</u>omain-Containing Protein 1 (TLNRD1) by the HUGO Gene Nomenclature Committee (Yates et al., 2017) reflecting its similarity to the R7R8 region of talin.

Talin1 and 2 are cytoplasmic adapters that provide a direct mechanosensitive link between the integrin family of cell adhesion molecules and the actin cytoskeleton (Calderwood et al., 2013; Goult et al., 2018). Talins are comprised of an N-terminal atypical FERM domain (Elliott et al., 2010), coupled via a flexible linker to a large rod domain comprised of 63 helices arranged into 13 helical bundles, R1-R13 (Fig.1A) (Goult et al., 2013; Gough and Goult, 2018). Twelve of the rod domains are arranged linearly, end-to-end to create the large extended talin rod domain which unfolds and stretches in response to mechanical force (Yao et al., 2016). However, the central region of the talin rod, comprising domains R7R8, adopts a unique fold, where R8, a 4-helix bundle, is inserted into a loop between two helices of the R7 5-helix bundle (Gingras et al., 2010) creating a branch in the talin rod (Fig. 1A). Our initial characterisation of R7R8 was facilitated by its sequence homology to TLNRD1; talin residues 1359-1659 show 22% identity to residues 44-352 of TLNRD1 (Fig. S1A) (Gingras et al., 2010), and this homology enabled us to pinpoint the nine helices that comprise talin R7R8, and facilitated resolution of the talin rod domain structure.

**Figure 1.**
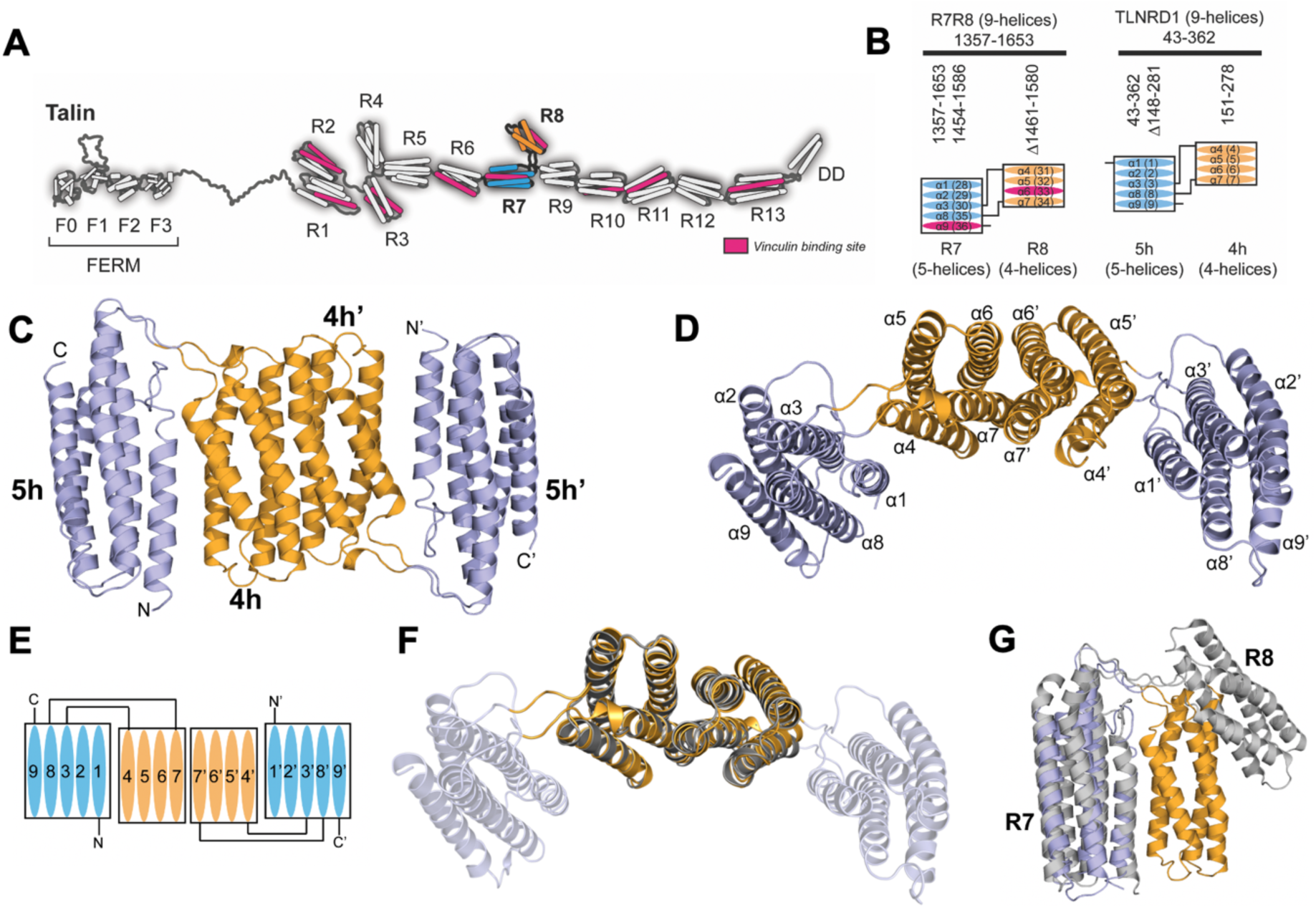
Talin Rod Domain Containing Protein 1 (TLNRD1) is homologous to talin R7R8. **(A)** The domain structure of talin. Talin has an N-terminal head domain (residues 1-400) comprising F0-F3 and a large rod domain comprised of 13 helical bundles, R1-R13 (Goult et al., 2013). The R7 (blue) and R8 (orange) domains are highlighted. The 11 vinculin binding sites (VBS) are shown in pink. **(B)** Schematic of the topology of R7R8 (left) and TLNRD1 (right) comprised of a 5-helix bundle (blue) with a 4-helix bundle (orange) inserted into the loop between helices α3 and α8. The VBS in R7 and R8 are highlighted, but are absent from TLNRD1 due to loss of the VBS consensus sequence (Gingras et al., 2010). **(C)** Crystal structure of TLNRD1-FL reveals an antiparallel, symmetric dimer. The domains of each monomer are labelled as 4h and 5h, and 4h’ and 5h’. **(D)** top down view of (C) with the helices labelled. The two 4-helix bundles dimerise via an extensive interface comprised of helices α6 and α7. **(E)** Schematic representation of the TLNRD1 dimer. **(F)** Overlay of the TLNRD1-FL and TLNRD-4H structures showing the dimerisation interface is the same in both. **(G)** Overlay of one monomer of TLNRD1 (orange and blue) with R7R8 (grey; PDB ID 2×0C (Gingras et al., 2010)). The individual domains are structurally homologous, but the orientation relative to each other is different, with talin R7R8 having a more open structure.

Talin domains R7R8 are emerging as a major signalling hub in talin, linking diverse cytoskeletal elements together (Goult et al., 2018). It forms part of a major actin-binding site in talin (termed ABS2) that spans R4-R8 (Atherton et al., 2015; Kumar et al., 2016; Hemmings et al., 1996). Interestingly, the unique topology of R7R8 positions R8 outside of the force transmission pathway in talin enabling it to remain folded in the presence of high mechanical forces (Yao et al., 2016; Yan et al., 2015). R8 has also been shown to bind to multiple other ligands, many of which contain Leucine-Aspartate motifs (LD-motifs) (Alam et al., 2014) including (i) Rap1-GTP-interacting adaptor molecule (RIAM), implicated in recruitment of talin to the leading edge of cells (Chang et al., 2014; Goult et al., 2013; Lee et al., 2009) and to filopodial protrusions (Lagarrigue et al., 2015) (ii) DLC1 (deleted in liver cancer 1), a RhoGAP that drives local modulation of actomyosin contractility (Zacharchenko et al., 2016; Li et al., 2011) and (iii) paxillin a focal adhesion adaptor protein implicated in numerous signalling pathways (Zacharchenko et al., 2016). R8 also contains a binding site for vinculin, a protein that crosslinks talin to actin and drives focal adhesion maturation (Yogesha et al., 2011; Gingras et al., 2010), and the intermediate filament protein α-synemin (Sun et al., 2008). The R7 domain also plays a major role in talin function, connecting adhesions to the microtubule network. Thus, R7 links to KANK proteins (Bouchet et al., 2016; Sun et al., 2016) which serve as platforms for the assembly of large Cortical Microtubule Stabilising Complexes that capture microtubules at the periphery of integrin adhesion complexes (Bouchet et al., 2016). This complex protein interaction network suggests that R7R8 plays a key role in coordinating multiple processes including crosstalk between all three cytoskeletal networks. Mutations in talin R7R8, obtained from the Catalogue of Somatic Mutations in Cancer (COSMIC), can perturb this coordination and increase invasion and migration in cells (Azizi et al., 2020).

Previous studies have shown that TLNRD1 is directly targeted by anti-oncogenic miRNAs, with an increase in TLNRD1 expression being associated with increased proliferation and xenograft growth in hepatocellular carcinoma. In contrast, TLNRD1 depletion reduced bladder cancer cell viability, migration and invasion (Tatarano et al., 2012; Wu et al., 2017). Here we show that although TLNRD1 and talin R7R8 both have the same domain structures and topology, and both bind actin, TLNRD1 is unique in its ability to bundle F-actin. Our structural data show that this is because it dimerizes via its 4-helix module. In U2OS cells, TLNRD1 localises to thick actin stress fibres and filopodial protrusions, with distinctive localisation to the filopodia tips. Overexpression of TLNRD1 also increases filopodia protrusion formation and cellular migration in 2D. Finally, we establish biochemically that TLNRD1 can interact with known R7R8 LD-motif containing ligands, potentially impacting their ability to interact with talin and therefore talin function in cells.

## Results

### The structural and sequence similarity of TLNRD1 and talin R7R8 suggest TLNRD1 arose through gene duplication

Based on sequence homology, TLNRD1 is predicted to have a structure similar to talin R7R8 with an additional unstructured region at the N-terminus (residues 1-43, Fig. 1B, S1A). The structural similarity of TLNRD1 with talin R7R8, plus their 22% amino acid sequence identity suggests that TLNRD1 may have originated from a gene duplication event. The *TLNRD1* gene first appears in choanoflagellates (*Monosiga brevicollis* and *Salpingoeca rosetta*) but is absent from earlier-diverging Opisthokont protists (e.g. *Abeoforma whisleri, Sphaeroforma arctica, Capsaspora owczarzaki*). This indicates that the gene duplication event that gave rise to TLNRD1 occurred around the time of the divergence of choanoflagellates from earlier Opisthokonts. A striking feature of the evolutionary history of TLNRD1 is that it is highly conserved throughout animal evolution, first appearing in sponges and choanoflagellates but absent in Cnidaria, nematodes, and arthropods (Fig. S1B).

While the R7R8 regions of the two mammalian talin isoforms are each encoded by 11 exons, the gene structure of TLNRD1 is markedly different. The entire mammalian *TLNRD1* gene consists of a single, large exon that encodes the whole protein. One possible explanation for the difference in gene structure between talin R7R8 and TLNRD1 is that a section of mature spliced talin mRNA encoding R7R8 was reintroduced back into an ancestral animal genome.

### Crystal Structure of TLNRD1

To determine the extent of the structural similarity of TLNRD1 with talin R7R8, we used X-ray crystallography to determine the structures of both the 4-helix domain (143-273, TLNRD1-4H) and the full-length protein (1-362, TLNRD1-FL). While the structure of TLNRD1-4H was straightforward to determine by molecular replacement with talin R7R8 (PDB: 2×0C (Gingras et al., 2010)), the structure of full length TLNRD1 was solved using the BALBES pipeline that performs molecular replacement using a repository of diverse structures from the PDB database (Long et al., 2008). TLNRD1-FL (PDB: 6XZ4) diffracted to 2.3 Å in *P*2_1_ space group with two TLNRD1 molecules in the asymmetric unit, and TLNRD1-4H (PDB: 6XZ3) to 2.2 Å in *I*4_1_22 space group with one molecule in the asymmetric unit (statistics in Table 1). In the case of full-length TLNRD1, electron density could only be traced for residues 40-341, whereas the 4-helix structure successfully resolved residues 148-270.

**Table 1.** Data collection and refinement statistics for TLNRD1 full-length and 4-helix domain.

| **Data collection** | **TLNRD1-FL** | **TLNRD1-4H** |
| --- | --- | --- |
| Synchrotron and Beamline | Soleil Proxima-1 | Soleil Proxima-1 |
| Space group | $P2_1$ | $I4_122$ |
| Molecule/a.s.u | 2 | 1 |
| Cell dimensions |  |  |
| $a, b, c$ (Å) | 69.17, 58.04, 84.32 | 114.59, 114.59, 59.40 |
| $\alpha, \beta, \gamma$ (°) | 90, 106.10, 90 | 90, 90, 90 |
| Resolution (Å) | 60.22 – 2.30<br>(2.38 – 2.30)* | 57.30 – 2.19<br>(2.31 – 2.19) |
| $R_{\text{merge}}$ | 0.087 (0.682) | 0.130 (1.034) |
| $I / \sigma I$ | 5.7 (1.3) | 12.5 (2.6) |
| $CC(1/2)$ | 0.994 (0.831) | 0.996 (0.938) |
| Completeness (%) | 98.4 (98.7) | 100 (99.9) |
| Redundancy | 2.8 (2.8) | 13.4 (13.6) |
| **Refinement** |  |  |
| Resolution (Å) | 2.30 | 2.19 |
| No. reflections | 27997 (2669) | 10464 (2428) |
| $R_{\text{work}} / R_{\text{free}}$ | 0.26/0.31 | 0.22/0.26 |
| No. atoms |  |  |
| Protein | 9047 | 1903 |
| Water | 70 | 38 |
| $B$-factors (Å<sup>2</sup>) | | |
| Protein | 74.25 | 72.05 |
| Water | 50.47 | 56.53 |
| R.m.s. deviations |  |  |
| Bond lengths (Å) | 0.004 | 0.003 |
| Bond angles (°) | 0.792 | 0.620 |
| Ramachandran plot |  |  |
| Favoured/allowed/<br>outlier (%) | 95.20/3.64/1.16 | 96.69/1.65/1.65 |
| Rotamer |  |  |
| Favoured/poor (%) | 88.31/4.18 | 96.12/0.97 |
| MolProbity scores |  |  |
| Protein geometry | 1.75 (97<sup>th</sup>)<sup>^</sup> | 1.01 (100<sup>th</sup>)<sup>^</sup> |
| Clash score all atoms | 1.88 (100<sup>th</sup>)<sup>^</sup> | 1.05 (100<sup>th</sup>)<sup>^</sup> |
| PDB accession no. | 6XZ4 | 6XZ3 |

The crystal structures confirmed that TLNRD1 consists of a 9-helix module comprised of a 4-helix and 5-helix bundle connected via the same unusual domain linkage identified in talin1 R7R8, where the 4-helix bundle is inserted into a loop between helices α3 and α4 of the 5-helix bundle (Fig. 1C-F). The first 40 residues of TLNRD1-FL are not visible in the density supporting the secondary structure prediction that this N-terminal region is predominantly disordered.

### TLNRD1 is a symmetric antiparallel dimer mediated by the 4-helix bundle

Our previous biochemical characterisation using gel filtration suggested that TLNRD1 runs as a dimer (Gingras et al., 2010). Here, the two TLNRD1 structures reveal the basis for TLNRD1 dimerisation which is mediated by a novel interface on the 4-helix bundle. In both the TLNRD1-FL and TLNRD1-4H structures, the 4-helix bundle forms an extensive interaction with a 4-helix bundle of a second TLNRD1 molecule (Fig. 1C). An identical dimer interface was observed in both the TLNRD1-FL and the TLNRD1-4H structures (Fig. 1F), which crystallised in different space groups (Table 1). Analysis of the macromolecular interface of the interaction using PISA (Krissinel and Henrick, 2007), verified that this was a *bona fide* dimer.

Dimerisation of TLNRD1 is mediated via an extensive hydrophobic interface on helices α6 and α7 with the two F250 side chains docking into the opposing molecule (Fig. 2A-B). The surface of each 4-helix bundle has a pocket created by F270’ and the small side chains of G217’ and A260’ that the F250 aromatic rings docks into (Fig. 2B). As it is a symmetric dimer, the F250’ docks into the equivalent pocket on the other molecule leading to an antiparallel configuration. The interface is capped at either end by electrostatic interactions between the side chains of E267 and R246’ and vice versa. Analysis of TLNRD1 over evolution from humans to the choanoflagellate, *Salpingoeca rosetta*, using the program ConSurf (Ashkenazy et al., 2016) reveals that these residues which mediate dimerisation are highly conserved (Fig. S2A) suggesting that TLNRD1 exists as an obligate dimer.

**Figure 2.**
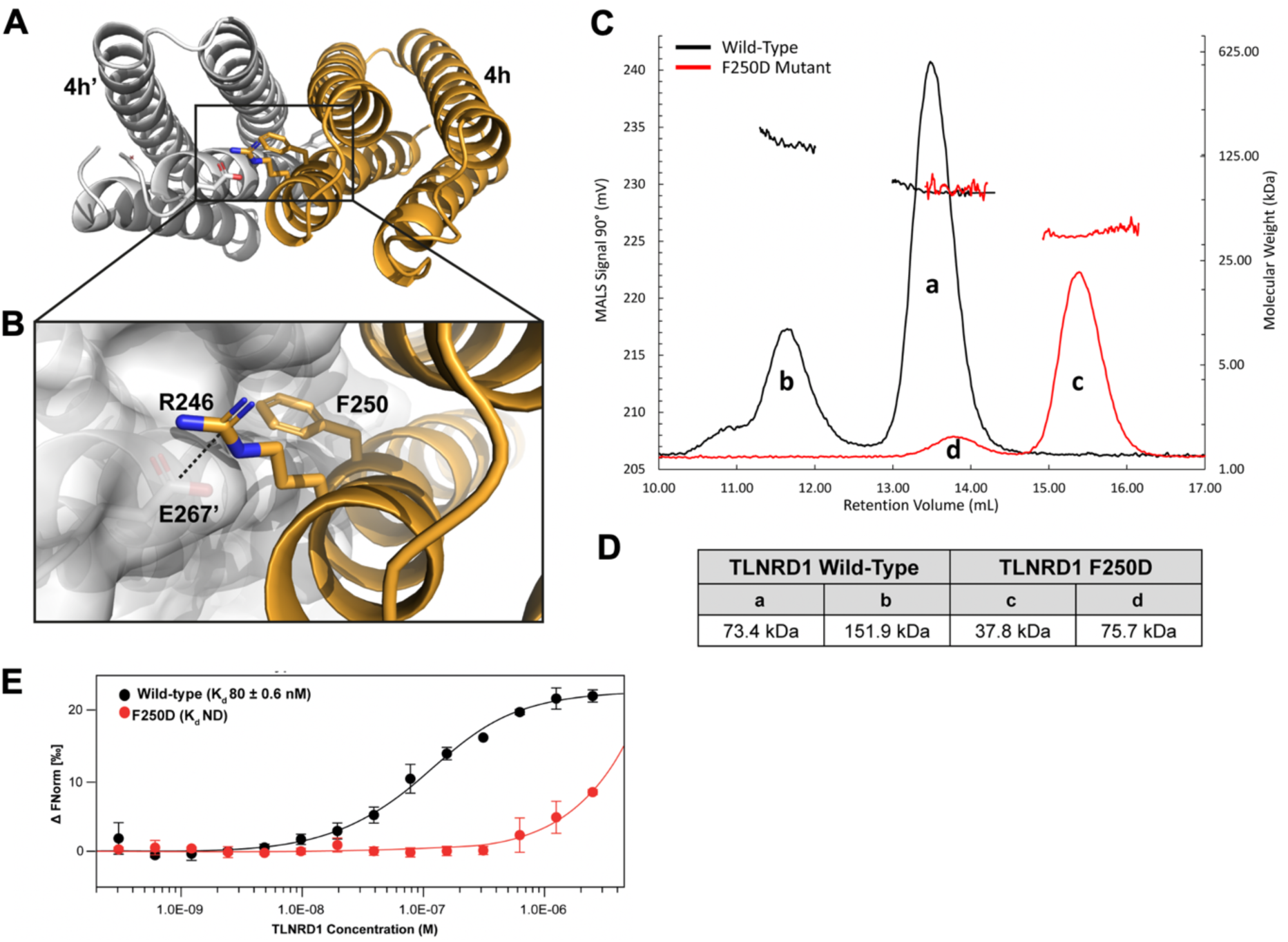
Characterisation of the TLNRD1 dimer. **(A-B)** The symmetrical antiparallel TLNRD1 dimer interface mediated by F250. **(B)** Zoomed in view. One monomer is shown as a surface representation, F250 from the opposing monomer (orange) inserts into the pocket on the surface of the other. A salt bridge between R246 and E267’ is shown. **(C-D)** SEC-MALS analysis of 106 μM TLNRD1-FL WT (black) and F250D (red). **(D)** Analysis of the molar mass of the major TLNRD1-FL species “a” yields a molecular weight of ∼74 kDa. The TLNRD1-F250D yields a molecular weight of ∼37.8kDa, peak “b”. **(E)** TLNRD1-FL dimerisation measured using MST at 25°C with 40% Red LED laser excitation. Unlabelled TLNRD1-FL was titrated into a fixed concentration (50 nM) of fluorescently labelled TLNRD1-FL. Data was analysed in MO.Affinity Analysis software (v2.1.3) using the law of mass action to generate a dimerisation K_d_ of 80 ± 0.59 nM. F250D K_d_ not determined.

We used size-exclusion chromatography multi-angle light scattering (SEC-MALS) to explore the oligomeric state of the TLNRD1 protein in solution. Both TLNRD1-FL (labelled “a” in Fig 2C) and TLNRD1-4H were dimeric at 25°C, and no monomer peak was present with either construct suggesting a high affinity dimer (Fig. 2C-D). TLNRD1-FL also showed a smaller tetramer peak (labelled “b” in Fig 2C), suggesting the presence of a dimer of dimers species. To explore the importance of F250 in dimerisation, we generated a point mutant of F250 that swapped the aromatic ring for a charged aspartate, F250D (TLNRD1-F250D). The TLNRD1-F250D mutant was a stable, folded, protein suitable for biochemical analyses (Fig. S2B), although the melting temperature, determined using Circular Dichroism, showed a significant reduction in thermostability (TLNRD1-FL T_m_ 69.7°C, TLNRD1-F250D T_m_ 48°C) (Fig. S2C). Analysis of the F250D mutant using SEC-MALS showed a clear transition from a dimeric to monomeric state (peak “c”), with only a small proportion remaining as a dimer (peak “d”), confirming the importance of this residue in mediating dimerisation (Fig. 2C-D).

The absence of a monomer peak on SEC-MALS suggests that the dimerisation of TLNRD1 is mediated by a high affinity interaction between F250 and the opposing pocket. We therefore used Microscale Thermophoresis (MST) to study the monomer-dimer equilibrium. In this experiment, unlabelled TLNRD1-FL was titrated against fluorescently-tagged TLNRD1-FL and a dimerisation constant (K_d_) of 80 nM obtained (Fig. 2E). MST confirmed that the F250D mutant prevented TLNRD1 dimerisation, as no K_d_ was generated. This high affinity suggests that TLNRD1 probably exists as an obligate dimer and a F250D mutant renders TLNRD1 monomeric.

### TLNRD1 has retained the functionality of talin R7R8

Talin R7R8 has emerged as a multifunctional signalling hub able to bind multiple ligands. Our previous studies on DLC1 binding to talin R8 (Zacharchenko et al., 2016; Li et al., 2011) and KANK proteins binding to talin R7 (Bouchet et al., 2016) have revealed that both R7 and R8 talin domains contain binding sites for proteins that contain LD-motifs (Alam et al., 2014). Similarly, RIAM and its paralogue lamellipodin have also been shown to interact with talin R8 via N-terminal LD-motifs (Goult et al., 2013; Chang et al., 2014). Furthermore, talin R7R8 also binds to actin filaments (Gingras et al., 2010) as part of actin-binding site 2 (Atherton et al., 2015; Kumar et al., 2016).

In light of this diverse R7R8 interactome and structural homology, we wanted to ask whether TLNRD1 is also able to bind to talin R7R8 ligands. We first tested RIAM and KANK1 as both contain well defined LD-motif talin binding sites, but RIAM binds to the 4-helix domain (R8 in talin), whereas KANK binds to the 5-helix domain (R7 in talin). Fluorescence polarisation studies showed that purified full-length TLNRD1 protein (residues 1-362, TLNRD1-FL, Fig 3A) interacts with RIAM TBS1 (residues 4-30) with a K_d_ of 0.25 µM (Fig. 3C), an interaction mediated by the 4-helix domain. Thus, the purified TLNRD1 4-helix domain alone (residues 143-273, TLNRD1-4H, Fig 3B) also bound to RIAM TBS1 with high affinity (K_d_ 0.59 µM) (Fig. 3D). Lamellipodin, a paralogue of RIAM, also interacts with TLNRD1-4H albeit with a lower K_d_ of 9 µM (Fig. 3D).

**Figure 3.**
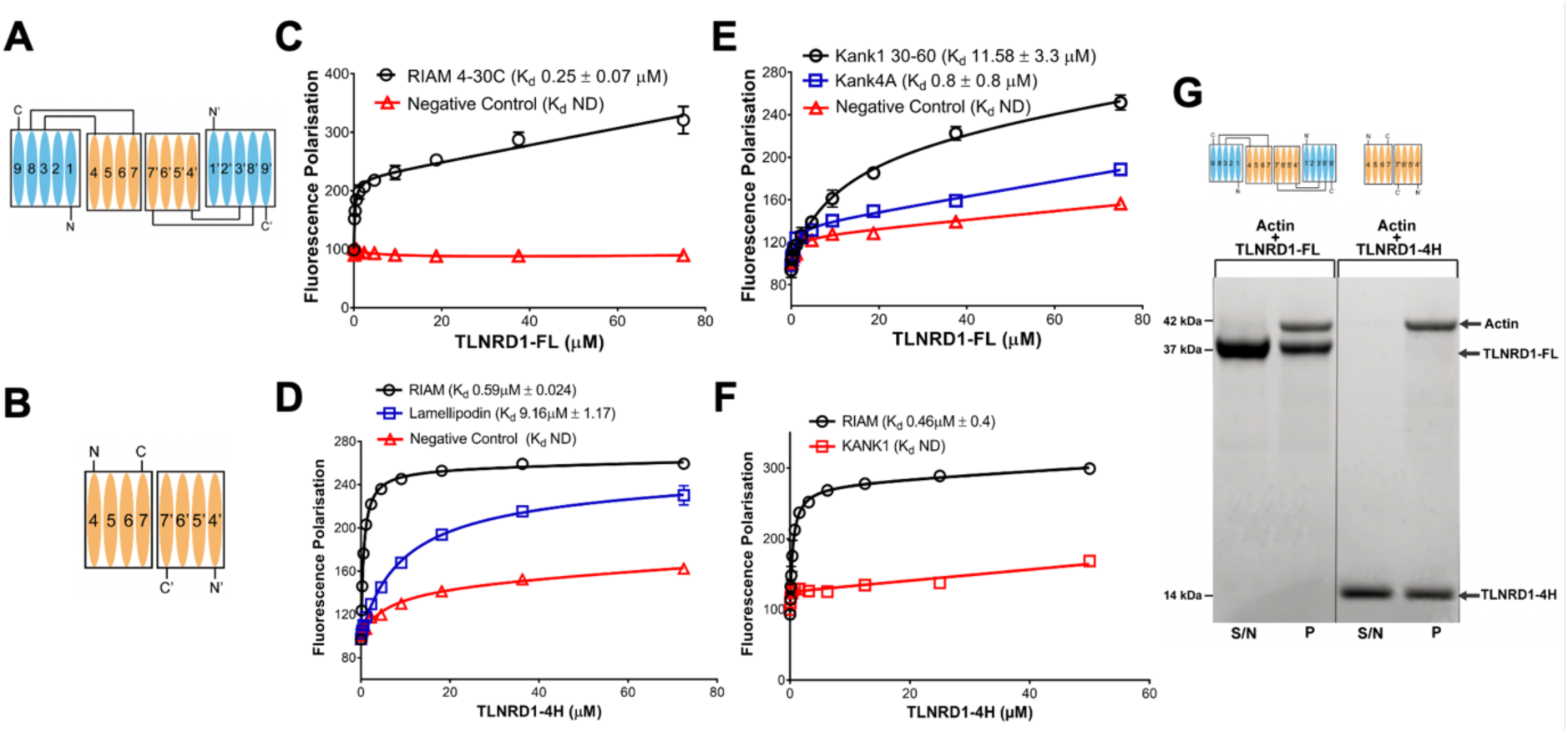
TLNRD1 has retained the ligand binding sites of talin R7R8. **(A-B)** Schematic of the TLNRD1-FL and TLNRD1-4H domain structures **(C-F)** TLNRD1 binds to LD-motif containing ligands. **(C-D)** Binding of fluorescein-labelled RIAM (residues 4-30) peptide with **(C)** TLNRD1-FL and **(D)** TLNRD1-4H measured using a Fluorescence Polarisation (FP) assay. The binding of Lamellipodin (residues 20-46; blue) is also shown**. (E-F)** Binding of BODIPY-labelled KANK1 (residues 30-60); wildtype (black) and 4A mutant (blue) peptides **(E)** binding to TLNRD1-FL and **(F)** not binding to TLNRD-4H, with RIAM (residues 4-30) positive control (black). ND = Not defined, experiments performed in triplicate. **(G)** High-speed actin cosedimentation assay showing that both TLNRD1-FL and TLNRD1-4H interact with F-actin.

The KANK1 “KN domain” LD-motif peptide (residues 30-60) also bound to TLNRD1-FL (K_d_ 11.6 µM) albeit with lower affinity than the 0.8 μM K_d_ with talin R7 (Fig. 3E). KANK1 binding to R7 was previously shown to require the LDLD sequence in the KN domain, with a quadruple mutation to alanine (4A mutant) abolishing binding to talin R7. Similarly, this KANK1 4A mutant showed a drastic reduction in binding to TLNRD1-FL confirming the interaction was LD-dependent. Lastly, no interaction was observed with KANK1 KN and TLNRD1-4H (Fig. 3F) confirming that the TLNRD1-KANK1 interaction is mediated via the 5-helix module in TLNRD1. Collectively, this biochemical analysis confirms that TLNRD1 has retained the LD-binding sites in both the 4-helix, and 5-helix modules after divergence from talin, and by extension that talin was able to bind LD-motifs prior to the duplication event.

Another major role of talin R7R8 is to bind actin, and based on this, TLNRD1 was previously shown to bind actin (Gingras et al., 2010). Using a high speed cosedimentation assay we confirmed that TLNRD1-FL interacts with actin filaments (Fig. 3G). Furthermore, we found that TLNRD1-4H alone is capable of interacting tightly with actin (Fig. 3G). This is surprising as, by itself, the equivalent 4-helix module in talin, R8, shows little actin binding in cosedimentation assays (Gingras et al., 2010), suggesting that TLNRD1 has enhanced actin filament binding. Taken together, these data indicate that TLNRD1 is an actin-binding protein that has retained functional similarities to talin R7R8.

### TLNRD1 is an Actin-Bundling Protein

TLNRD1, like talin R7R8, binds to actin in *in vitro* actin cosedimentation assays (Fig. 3G; (Gingras et al., 2010)). In talin, the second actin-binding site (ABS2) maps to domains R4-R8 in the talin rod, with the two 4-helix bundles, R4 and R8 engaging the actin filament (Fig. 4A) (Atherton et al., 2015; Kumar et al., 2016). The actin-binding surface on R8 maps onto one face (helices α2 and α3) of the bundle. Our initial hypothesis was that as TLNRD1 was a dimer it would also have two 4-helix domains that together might engage a single actin filament in a similar fashion to talin ABS2. However, the structure of TLNRD1 dimer shows that the putative actin-binding surface on α2 and α3 is positioned facing out from the dimer interface (Fig. 4A). This arrangement raised the possibility that TLNRD1 might be able to engage two actin filaments simultaneously, and thus crosslink and bundle actin.

**Figure 4.**
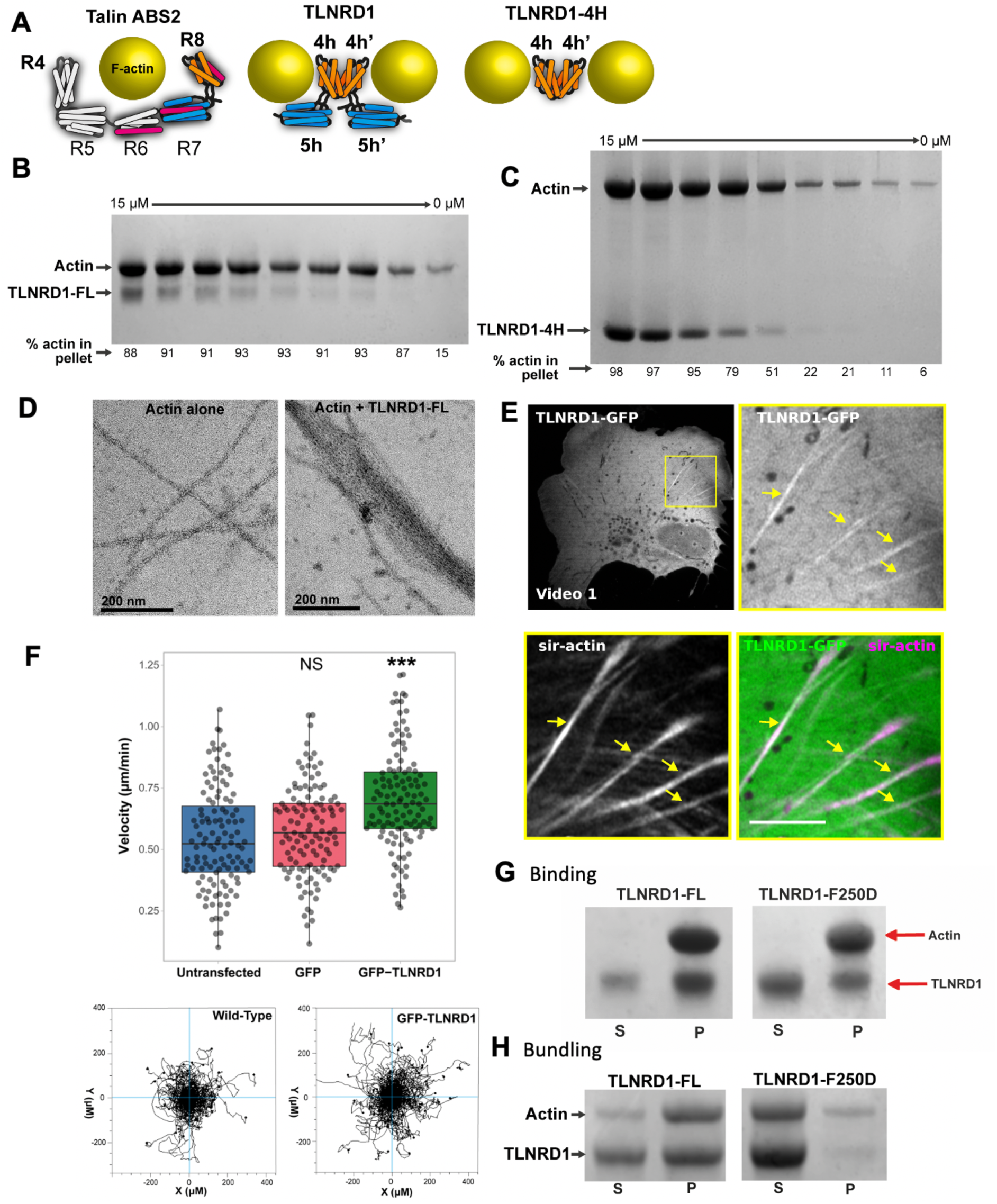
TLNRD1 is an actin-bundling protein. **(A)** Schematic of the talin actin-binding site 2 (ABS; domains R4-R8) binding a single actin filament (left), TLNRD1-FL (middle) and TLNRD1-4H (right) binding to two actin filaments. **(B-C)** Low-speed actin cosedimentation bundling assay with TLNRD1-FL **(B)** and TLNRD1-4H **(C)** serial dilution against 25 µM F-actin. **(D)** Transmission electron micrographs of negatively stained F-actin alone (left) and with TLNRD1-FL showing bundled actin filaments (right). **(E)** U2OS cells expressing GFP-TLNRD1 were plated on fibronectin, incubated for 2 hr with SiR-actin to label the actin cytoskeleton, and imaged live using an Airyscan confocal microscope (1 picture every 16 s; Video S1). Scale bars: (inset) 5 mm. The yellow arrows highlight GFP-TLNRD1 localising to actin stress fibres. **(F)** Random 2D migration assay of U2OS cells plated on fibronectin and non-transfected or transiently expressing GFP or GFP-TLNRD1. GFP-TLNRD1 overexpression increases migration velocity in 2D (n = 120 cells from 3 independent repeats, ***p value <0.001). Cell trajectories for non-transfected and GFP-TLNRD1 expressing cells are shown. **(G)** High speed actin binding assay of TLNRD1-FL and TLNRD1-F250D showing little impact on actin binding with loss of dimerisation. **(H)** Low speed actin cosedimentation assay showing loss of TLNRD1 actin bundling with loss of dimerisation as a result of the F250D mutation.

To establish whether TLNRD1 is also an actin-bundling protein, we used a low speed (10k rpm) cosedimentation assay where actin filaments remain in solution and only bundles of actin filaments will sediment with bundling proteins. At low speed, actin by itself remained in the supernatant, but addition of TLNRD1-FL resulted in a clear increase in the levels of actin in the pellet confirming that TLNRD1 is an actin bundler (Fig. 4B). In contrast neither talin R7R8, nor ABS2, which are both monomeric are able to bundle actin (Atherton et al., 2015). Following on from the discovery that the 4-helix domain of TLNRD1 alone can bind actin filaments (Fig. 3G), we tested the ability of TLNRD1-4H to bundle actin in the low speed actin cosedimentation assay and surprisingly we found that the 4-helix domain by itself, is an effective actin bundler (Fig. 4C). To confirm TLNRD1 bundling activity, actin filaments were visualised by negative stain electron microscopy both in the absence, or presence, of TLNRD1-FL (Fig. 4D). This revealed that TLNRD1-FL can form large bundles of actin filaments with tight inter-filament spacing. In addition, TLNRD1 was found to decorate actin stress fibres when expressed as N-terminally GFP-tagged (GFP-TLNRD1) in U2OS cells (Fig. 4E). The effect of TLNRD1 on cell migration was assessed using live-cell imaging of U2OS cells plated on fibronectin. Tracking cell migration over time revealed that expression of GFP-TLNRD1 significantly increased migration velocity compared to non-transfected, or GFP expressing, cells (Fig. 4F).

A key feature which distinguishes TLNRD1 from R7R8 is its dimeric state. To explore the importance of TLNRD1 dimerisation in promoting bundling of actin filaments, actin cosedimentation assays were performed with the TLNRD1-F250D dimerisation mutant. TLNRD1-F250D was still able to bind to actin filaments (Fig. 4G), however, it showed significantly reduced actin-bundling activity (Fig. 4H), confirming that dimerisation is required for TLNRD1 to bundle actin. As TLNRD1 binds actin considerably more tightly than talin R7R8, as well as talin ABS2, and can also drive actin bundling, its interactions with actin are distinct from the established talin ABS2 interaction.

### TLNRD1 promotes filopodia formation and co-localises with myosin-X at filopodia tips

As actin-bundling proteins often contribute to filopodia functions (Gupton and Gertler, 2007; Khurana and George, 2011; Jacquemet et al., 2015), we next investigated whether TLNRD1 might also modulate filopodia formation. U2OS cells were transfected with MYO10-mScarlet (to induce and visualize filopodia) together with GFP, GFP-TLNRD1 or GFP-TLNRD1-F250D, imaged on a spinning disk confocal microscope, and the number of MYO10-positive filopodia was scored (Fig. 5A-B). Over-expression of GFP-TLNRD1 promoted filopodia formation whereas the expression of GFP-TLNRD1-F250D did not, indicating that TLNRD1 dimerisation is required to promote filopodia formation (Fig. 5A-5B). To further validate a role for TLNRD1 in modulating filopodia formation, TLNRD1 expression was silenced in U2OS cells using two independent small interfering RNAs (siRNAs) (Fig. 5C-D). TLNRD1 gene silencing efficiency could be validated using qPCR (Fig. 5C) and western blotting following immuno-precipitation (Fig. 5D). Silencing of TLNRD1 led to a decrease in the number of MYO10 filopodia in U2OS cells (Fig. 5E). Interestingly, when imaging GFP-TLNRD1 together with MYO10, TLNRD1 appeared to localise to filopodia (Fig. 5A). To gain insights into the spatial distribution of TLNRD1 in filopodia, U2OS cells transiently expressing mScarlet-MYO10 and GFP-TLNRD1 or GFP-TLNRD1-F250D were imaged using structured-illumination microscopy (SIM) (Fig. 5F). The average distribution of these two TLNRD1 constructs in filopodia was then mapped as previously described (Fig. 5G) (Jacquemet et al., 2019). Surprisingly, these experiments revealed that TLNRD1, unlike other filopodia bundling proteins such as fascin, accumulates to filopodia tips. Altogether, our data demonstrates that TLNRD1 is a filopodia tip protein which modulates filopodia formation.

**Figure 5.**
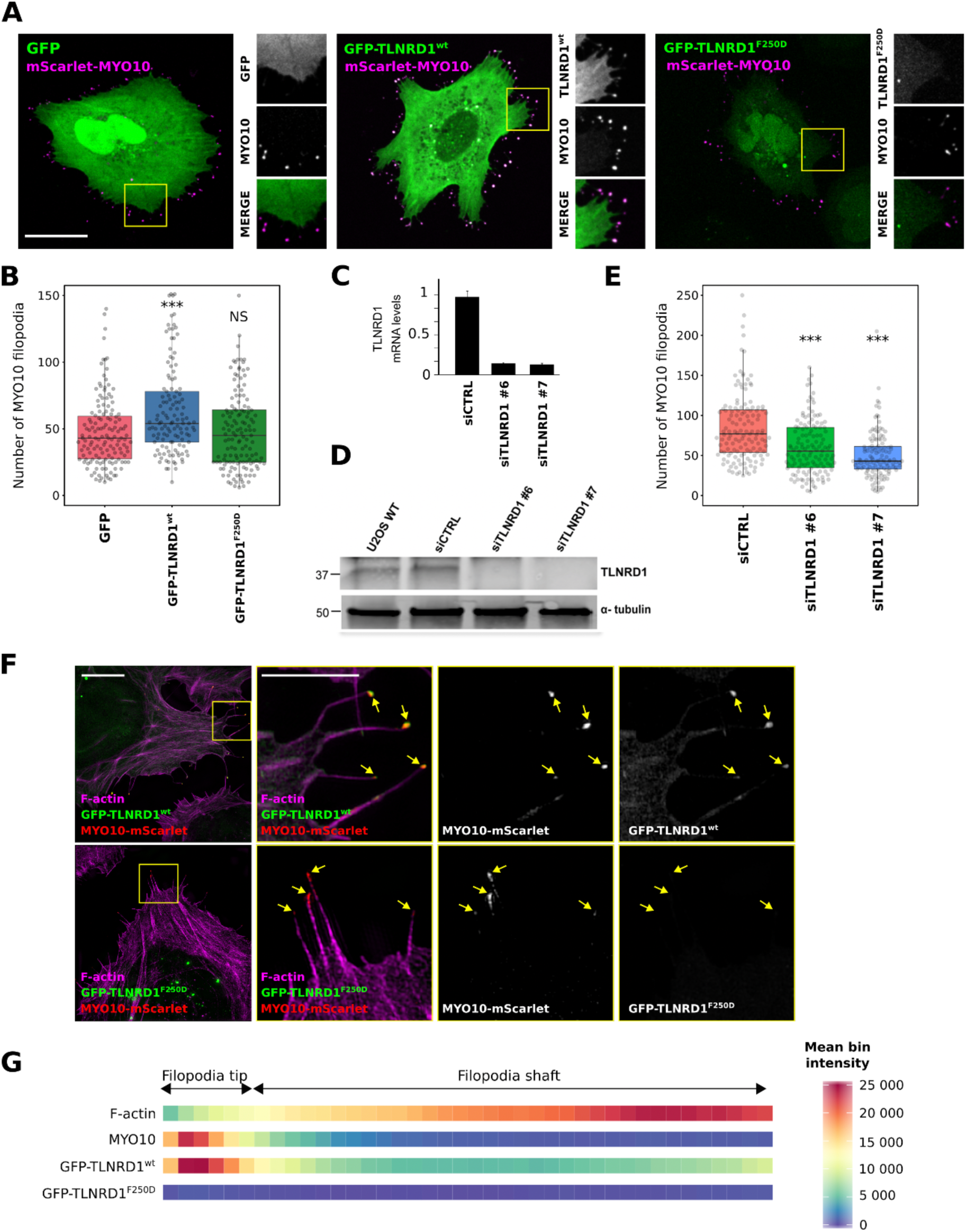
TLNRD1 localises to, and modulates, filopodia. **(A-B)** U2OS cells transiently expressing MYO10-mScarlet and GFP, GFP-TLNRD1 or GFP-TLNRD1-F250D were plated on fibronectin for 2 hr, fixed, imaged using a spinning disk confocal microscope. Representative images are displayed (**A**). Yellow square highlights region of interest (ROI) which is magnified. Scale bar: 25 μm. **(B)** The number of MYO10-positive filopodia per cell was then quantified (n > 125 cells, three biological repeats; *** p value = 0.002). **(C-D)** Efficiency of siRNA-mediated (oligos nos. 6 and 7) silencing of TLNRD1 in U2OS cells as detected using qPCR **(C)** or western blotting **(D)**. To reliably detect TLNRD1 protein levels by western blot, TLNRD1 was first immuno-precipitated from cell lysate as indicated. (**E**) TLNRD1-silenced (oligos nos. 6 and 7) U2OS cells transiently expressing MYO10-GFP were plated on fibronectin for 2 hr, fixed, and the number of MYO10-positive filopodia per cell was quantified (n >150 cells, three biological repeats; ***p value <0.001). **(F-G)** U2OS cells expressing MYO10-mScarlet and GFP-TLNRD1 or GFP-TLNRD1-F250D were plated on fibronectin for 2 hr; stained for F-actin and imaged using SIM. (**F**) Representative maximum intensity projections are displayed. The yellow squares highlight ROIs, which are magnified; scale bars: (main) 10 μm; (inset) 5 μm. (**G**) Heatmap highlighting the subcellular localization of F-actin, MYO10, TLNRD1 and TLNRD1-F250D within filopodia based on >360 intensity profiles (see methods for details). P-values were determined using a randomization test (see methods for details).

## Discussion

Talin Rod Domain Containing Protein 1 (TLNRD1) shares high sequence and structural similarity with the R7R8 region of talin, an important signalling hub which coordinates multiple cellular pathways. In this work we have characterised TLNRD1 as a novel actin-bundling protein that supports the formation of filopodia.

Talin R7R8 has been shown to have multifunctional roles; binding to actin and coupling this to multiple ligands containing LD-motif sequences. Here we show that TLNRD1 has retained R7R8’s diverse interactions through its capacity to bind LD-motifs and actin filaments. With the high similarity between talin R7R8 and TLNRD1, it is possible that TLNRD1 might act as a dominant negative modulator of talin function, fine-tuning talin signalling responses by sequestering R7R8 ligands and thus uncoupling them from their connection to integrin-mediated adhesion complexes. TLNRD1 binding to LD-motif containing ligands is expected to be more diffusive as R7R8 is a part of the talin rod and is more restricted in its mobility as it is tethered in adhesive structures. TLNRD1 on the other hand offers the same potential functionality but without the key domains of talin which coordinate its adhesion complex behaviour.

Our data also demonstrate that TLNRD1 is an obligate dimer with a mode of actin binding distinct from talin ABS2. The orientation of the two TLNRD1 monomers with respect to actin allows the dimeric protein to act as an efficient actin bundler. Together, this suggests that TLNRD1 can perform similar, as well as distinct, functions to talin R7R8.

### Compact arrangement of the bundles

Another notable feature of the TLNRD1 structure is that the 5-helix bundle packs against the side of the 4-helix bundle. In contrast, multiple structures of talin1 R7R8 have been solved alone (Gingras et al., 2010), and in complex with DLC1 (Zacharchenko et al., 2016) and RIAM (Chang et al., 2014), and in all of these structures, there are no interactions between the R7 and R8 domains, with R8 protruding away from R7.

In both the TLNRD1 and R7R8 structures, the linker connecting the two domains forms a two-stranded anti-parallel ß-sheet with complementary backbone hydrogen bonding (Gln-147 bonds with Ala-282 and Gln-152 with Arg-271 in TLNRD1) holding the two linkers in close proximity. This ß-sheet arrangement means there is limited flexibility between the two domains. However, the structures reveal that there is sufficient flexibility to allow both an open conformation, as in the talin R7R8 structures, and a more compact arrangement as in the TLNRD1 structure. It will be interesting to determine whether opening and closing of the two domains relative to each other is of biological relevance; it is tempting to speculate that a closed conformation could represent an autoinhibited conformation rendering some binding sites cryptic. However, the limited specific inter-domain contacts in the TLNRD1 structure suggests that the more compact structure may simply represent tight packing as a result of crystal formation, although it is interesting to note that we observe a similar compact conformation in solution using Small Angle X-Ray Scattering (SAXS) (Fig. S3).

### The role of TLNRD1 in filopodia functions

We report here that TLNRD1 localises to and regulates filopodia formation (Figure 5). While the precise mechanism by which TLNRD1 induces filopodia remains to be investigated, we found that TLNRD1 dimerisation is required. As TLNRD1 dimers can bundle actin and as actin bundlers have a well-characterised role in inducing filopodia (Gupton and Gertler, 2007; Khurana and George, 2011; Jacquemet et al., 2015) it is tempting to speculate that TLNRD1 promotes filopodia formation via its actin-bundling property. Interestingly, previously described filopodia actin-bundling proteins, such as fascin, typically localise to filopodium shafts (Pfisterer et al., 2020) where they tightly pack parallel actin filaments into bundles which allow filopodia structures to stabilise (Gupton and Gertler, 2007; Khurana and George, 2011; Jacquemet et al., 2015). In particular, the interaction between fascin and actin has been recently demonstrated to be the highest at filopodia base (Pfisterer et al., 2020). In contrast, TLNRD1 accumulated almost exclusively to filopodium tips and the contributions of an actin bundler to filopodia tip functions remains to be studied.

How TLNRD1 localises to filopodia tips remains unknown, but we speculate that it is via a direct interaction with a filopodia tip complex protein. For instance, we demonstrated here that TLNRD1 could interact directly in vitro with lamellipodin, which also localises to filopodia tips (Krause et al., 2004). Interestingly, TLNRD1 localisation within filopodia is very similar to that of talin, which also accumulate at filopodia tips (Jacquemet et al., 2019; Lagarrigue et al., 2015; Jacquemet et al., 2016). We previously showed that talin regulates filopodia formation by tightly coordinating integrin activity within filopodia (Jacquemet et al., 2016; Miihkinen et al., 2020). Therefore, future work will aim at investigating the role of TLNRD1 in modulating integrin activity in cells and in filopodia.

### Under the radar

Inspection of the gene expression in the EBI Expression Atlas (Petryszak et al., 2016) shows that TLNRD1 is expressed across most tissues throughout embryonic stages and into adulthood, suggesting a widespread, cellular function. Yet prior to this work, only a handful of papers have looked at this protein. The original name MESDC1 originated from its identification as a candidate essential gene for mesoderm development (Wines et al., 2001), a role that was later attributed to MESDC2 (Hsieh et al., 2003). Our 2010 study on talin R7R8 provided the initial characterisation of the MESDC1 protein (Gingras et al., 2010) which prompted its renaming to TLNRD1. Since then, TLNRD1 has been implicated as a putative oncogene regulated by microRNAs (Wu et al., 2017; Tatarano et al., 2012).

It is possible to speculate that the lack of focus on TLNRD1 arose due to the confluence of several factors. The absence of any reported diseases arising from mutations in TLNRD1 meant it escaped medical attention, furthermore, TLNRD1 has a unique evolutionary history, showing high conservation throughout vertebrate evolution with early homologs in sponges and choanoflagellates but has been lost in some key model organisms including *Drosophila*, yeast and *C. elegans*. The absence from the model organisms, *Drosophila*, C.elegans and yeast meant it was not picked up in genetic screens. Lastly, until recently the only commercially available anti-MESDC1 antibody (Abcam) did not recognise endogenous TLNRD1. Unfortunately, we could not study the localisation of endogenous TLNRD1 in cells as none of the antibodies we tested worked for immunofluorescence.

### The role of TLNRD1 in cancer

Two studies on potential oncogenes regulated by anti-oncogenic micro-RNAs; miR-508-5p in hepatocellular carcinoma (Wu et al., 2017) and miR-574-3p in bladder cancer (Tatarano et al., 2012), have both implicated TLNRD1 as a novel oncogene. Interestingly data from multiple databases (Nagy et al., 2018) shows that TLNRD1 mRNA levels often correlate with poor lung cancer patients survival. Our data on over-expression of TLNRD1 in osteosarcoma cells indicate that TLNRD1 over-expression drives cell migration. In addition, the number of filopodia per cell is significantly increased when TLNRD1 levels are elevated which may drive a more invasive morphology (Jacquemet et al., 2017). Future work will aim at elucidating the contribution of TLNRD1 to invasive cell migration.

In summary, we have structurally and biochemically characterised the TLNRD1 protein, as a novel actin-bundling protein that can drive filopodia formation and cell migration. In doing so we have expanded both the talin family of proteins with the identification of a third talin gene and also the growing set of proteins that can polymerise actin filaments into bundles.

## Materials and Methods

### Constructs for Biochemical/Structural Assays

Human TLNRD1 (TLNRD1-FL, residues 1-362) pET151 was purchased as a codon optimised synthetic gene from GeneArt (Regensburg, Germany). TLNRD1 4-helix domain (TLNRD1-4H, residues 143-273) was sub-cloned into pET151 vector. Single point mutations were introduced into TLNRD1 using site directed mutagenesis with *Pfu* DNA polymerase (Promega, Madison US), followed by digestion with DpnI at 37°C for 1 hr and transformation into DH10β *E. coli* cells.

All expression constructs have been deposited in Addgene www.addgene.org/ben_goult

### Protein Expression and Purification

TLNRD1 constructs were expressed in BL21(DE3) *E. coli* cells grown in Lysogeny broth (LB) at 37°C with 100 μg/ml ampicillin. Expression was induced with 0.1 mM Isopropyl β-D-1-thiogalactopyranoside (IPTG) and cells further incubated at 18°C overnight. Following centrifugation, pelleted cells were resuspended in nickel affinity buffer (50 mM imidazole, 500 mM NaCl, 20 mM Tris-HCl, pH 8). Cell lysates were loaded onto a 5 ml HisTrap HP column (GE Healthcare) for purification by nickel affinity chromatography, as described previously (Khan et al., 2020). Eluted protein was exchanged into MES buffer (20 mM 2-N-morpholinoethanesulfonic acid, 20 mM NaCl, 2 mM DTT, pH 6.5). His-tags were removed with AcTEV protease (Invitrogen) overnight and proteins further purified with a HiTrap SP HP cation exchange column (GE Healthcare).

### Fluorescence Polarisation Assay

Assays were performed with a serial dilution of protein, with target peptides at a concentration of 100 nM. The peptides, synthesised by GLBiochem (China), were coupled with either a fluorescein or BODIPY-TMR dye (ThermoFisher). Fluorescence polarisation was measured using a CLARIOstar plate reader (BMGLabTech) at 20°C. Data was analysed using GraphPad Prism 7 software and K_d_ values generated with the one-site total binding equation.

### F-actin co-sedimentation Assays

Actin was isolated from rabbit muscle acetone powder kindly gifted by Professor Mike Geeves using a previously described protocol (Spudich and Watt, 1971). Final purified F-actin was stored at 4°C in polymerisation buffer (10 mM Tris-HCl pH 7, 50 mM NaCl, 2 mM MgCl_2_, 1 mM NaN_3_, 1 mM DTT). For cosedimentation assays, F-actin was diluted to 15 µM and incubated with a serial dilution of protein starting at a 1:1 ratio for 1 hr at room temperature. To test binding, samples were spun at 100,000 x g for 20 min at 4°C. To test bundling activity, samples were spun at 10,000 x g for 15 min at 4°C. Equal volumes of pellet and supernatant were loaded onto SDS-PAGE gels and densities analysed using ImageJ software (Schneider et al., 2012).

### Electron Microscopy

F-actin was diluted to 25 µM in polymerisation buffer. TLNRD1-FL was incubated with the F-actin at a 1:1 ratio at room temperature for 1 hr. After incubation, samples were diluted down to 2 µM with polymerisation buffer. Samples were applied to 300 mesh carbon-coated copper grids for 30 seconds, excess solution removed and negatively stained for 1 minute with 2% (wt/vol) uranyl acetate. Excess stain was removed and grids washed under a stream of ddH_2_O. Grid samples were dried before imaging. Images were taken on a JEOL-1230 transmission electron microscope equipped with a Gatan One View 16 MP camera and an accelerating voltage of 80 kV.

### Size Exclusion Chromatography Multi-Angle Light Scattering (SEC-MALS)

SEC-MALS analysis of TLNRD1-FL and TLNRD1-F250D mutant was performed at room temperature with 100 μl of protein at 150 μM. Samples were loaded onto a Superdex 75 size-exclusion column (GE Healthcare Life Sciences) and eluted proteins measured by Viscotek Sec-Mals 9 and Viscotek RI detector VE3580 (Malvern Panalytical). Data was analysed using OmniSEC software.

### Crystallisation, X-Ray Data Collection and structure solution

TLNRD1-FL and TLNRD1-4H crystallisation trials were performed using hanging drop vapour diffusion at 21°C with concentrations of 390 µM and 350 µM respectively. Crystals of TLNRD1-FL were obtained in a condition containing 250 mM NaSCN, 20% PEG3350 and grown to optimal size in 7 days. For TLNRD1-4H, crystals were obtained in 300 mM HOC(CO_2_H)(CH_2_CO_2_NH_4_)_2_, 25% PEG3350 and attained full growth in 4 days. Crystals were harvested in their respective growth solutions supplemented with 25% ethylene glycol as cryoprotectant, mounted on CryoLoops (Hampton research) or LithoLoops (Molecular Dimensions) and vitrified in liquid nitrogen for data collection. X-ray diffraction data sets were collected at 100 K at Proxima-1 beamline at Soleil synchrotron (Paris, France) using a Pilatus 6M detector (Dectris, Baden, Switzerland) and processed by autoPROC pipeline (Vonrhein et al., 2011) which incorporates XDS (Kabsch, 2010), AIMLESS (Evans and Murshudov, 2013) and TRUNCATE (Evans, 2011) for data integration, scaling and merging respectively. The structure of TLNRD1-4H was determined by molecular replacement carried out by PHASER (McCoy et al., 2007) using 2×0C as search model. For the TLNRD1-FL solution BALBES molecular replacement pipeline (Long et al., 2008) was employed to generate the initial model which was then manually tweaked prior to adjustment and refinement. Manual model adjustment was carried out in COOT (Emsley et al., 2010) followed by refinement using PHENIX.REFINE (Afonine et al., 2012). Interaction properties of the dimer interface of the TLNRD1-FL were assessed by PISA (Krissinel and Henrick, 2007) and figures were prepared in PyMOL (Schrödinger LLC, Cambridge MA, USA). Models were validated by MOLPROBITY (Chen et al., 2010) prior to deposition.

### Size Exclusion Chromatograph-Small Angle X-ray Scattering (SEC-SAXS)

SEC-SAXS data was collected at Diamond Light Source beamline B21 (Didcot, UK). TLNRD1-FL SAXS experiments were performed at 80 µM and 185 µM in 20 mM Tris pH 7, 50 mM NaCl, 2 mM DTT. All samples were loaded onto a KW-403-4F 10-600 kDa size-exclusion column (Shodex) connected to an Agilent 1200 HPLC system. Data was analysed using ScÅtter software available from http://www.bioisis.net/ and ATSAS online services (Franke et al., 2017).

### Microscale Thermophoresis

For investigations into dimerisation, his-tagged TLNRD1 was diluted to 100 nM and coupled with His-tag NT-647 dye (RED-tris-NTA NanoTemper, München, Germany) at room temperature for 30 min. Unlabelled non-his-tagged TLNRD1 was diluted down to 5 µM. Final working concentrations of labelled and unlabelled protein were 50 nM and 2.5 µM respectively with the unlabelled protein serially diluted. Samples were loaded into Monolith NT.115 Capillaries and ran on a Monolith NT.115 (NanoTemper, München, Germany). All experiments were run at 25°C with 40% laser excitation. Data was analysed using MO.Affinity Analysis v2.3.

### Cell Culture

For cell culture experiments, N-terminal GFP tagged mouse TLNRD1 was used as previously described (Gingras et al., 2010). The mScarlet-MYO10 construct was described previously (Jacquemet et al., 2019). Human U2OS osteosarcoma cells (Leibniz Institute DSMZ-German Collection of Microorganisms and Cell Cultures, Braunschweig DE) were grown in DMEM supplemented with 10% FCS, 2 mM L-glutamine, 1% (vol/vol) penicillin and streptomycin, and maintained at 37°C in a humidified 5% CO_2_ environment.

### Transfection and siRNA knockdown

Plasmids of interest were transfected using Lipofectamine 3000 and the P3000TM Enhancer Reagent (ThermoFisher Scientific) according to the manufacturer’s instructions. The expression of proteins of interest was suppressed using 100 nM siRNA and lipofectamine 3000 (ThermoFisher Scientific) according to manufacturer’s instructions. The siRNA used as control (siCTRL) was Allstars negative control siRNA (QIAGEN, Cat. No. 1027281). The siRNAs targeting TLNRD1 were purchased from QIAGEN (siTLNRD1#6, Hs_MESDC1_6 FlexiTube siRNA, Cat. No. SI04217605; siTLNRD1 #7, Hs_MESDC1_7 FlexiTube siRNA, Cat. No. SI04314569, siTLNRD1#8, Hs_MESDC1_8 FlexiTube siRNA, Cat. No. SI04362820).

### TLNRD1 Immunoprecipitation

For immunoprecipitation experiments, U2OS cells were grown to 90% confluency in a 100 mm dish and lysed with equal volume of appropriate buffer (40 mM HEPES-NaOH pH 7.4, 75 mM NaCl, 2 mM EDTA, 2% NP40). Lysates were cleaned by centrifugation at 10,000 x g for 5 minutes at 4°C before 3 hr incubation at 4°C with Dynabeads Protein G superparamagnetic beads pre-coated with anti-TLNRD1 antibody or IgG control. Beads were washed three times with PBS. Protein extracts were separated under denaturing conditions by SDS–PAGE and subjected to western blotting with appropriate primary antibody diluted 1:1000 followed by incubation with the appropriate fluorophore-conjugated secondary antibody diluted 1:5000. Membranes were scanned using an Odyssey infrared imaging system (LI-COR Biosciences).

Anti-TLNRD1 antibodies were raised in rabbit against recombinantly expressed human TLNRD1 (residues 1-362) by Capra Science (Sweden). The secondary antibody used for western blot was an IRDye 800CW conjugated donkey anti-rabbit antibody (Li-Cor, cat number 926-32213).

### Sample preparation for light microscopy

For SIM imaging, U2OS cells transiently expressing GFP-TLNRD1 and Myosin-X-mScarlet were plated on high tolerance glass-bottom dishes (MatTek Corporation, coverslip #1.7) pre-coated first with Poly-L lysine (10 mg/ml, 1 hr at 37°C) and then with bovine plasma fibronectin (10 mg/ml, 2 hr at 37°C). After 2 hr, samples were fixed and permeabilised simultaneously using a solution of 4% (wt/vol) PFA and 0.25% (vol/vol) Triton X-100 for 10 min. Cells were then washed with PBS, quenched using a solution of 1 M glycine for 30 min, and incubated with SiR-actin (100 nM in PBS; Cytoskeleton; catalogue number: CY-SC001) at 4°C until imaging (minimum length of staining, overnight at 4°C; maximum length, 1 week). Just before imaging, samples were washed three times in PBS and mounted in vectashield (Vectorlabs).

To map the localization of each protein within filopodia, images were first processed in Fiji (Schindelin et al., 2012) and data analysed using R as previously described (Jacquemet et al., 2019). Briefly, in Fiji, the brightness and contrast of each image was automatically adjusted using, as an upper maximum, the brightest cellular structure labelled in the field of view. In Fiji, line intensity profiles (1 pixel width) were manually drawn from filopodium tip to base (defined by the intersection of the filopodium and the lamellipodium). To avoid any bias in the analysis, the intensity profile lines were drawn from a merged image. All visible filopodia in each image were analysed and exported for further analysis (export was performed using the ‘‘Multi Plot’’ function). For each staining, line intensity profiles were then compiled and analysed in R. To homogenize filopodia length, each line intensity profile was binned into 40 bins (using the median value of pixels in each bin and the R function ‘‘tapply’’). Using the line intensity profiles, the percentage of filopodia positive for each POI was quantified. The map of each POI was created by averaging hundreds of binned intensity profiles.

For the filopodia formation assays, cells were plated on fibronectin-coated glass-bottom dishes (MatTek Corporation) for 2 hr. Samples were fixed for 10 min using a solution of 4% (wt/vol) PFA, then permeabilised using a solution of 0.25% (vol/vol) Triton X-100 for 3 min. Cells were then washed with PBS and quenched using a solution of 1 M glycine for 30 min. Samples were then washed three times in PBS and stored in PBS containing SIR-actin (100 nM; Cytoskeleton; catalogue number: CY-SC001) at 4°C until imaging. Just before imaging, samples were washed three times in PBS. Images were acquired using a spinning disk confocal microscope (100x objective). The number of filopodia per cell and their length was manually scored using Fiji.

### Microscopy setup

The spinning disk confocal microscope used was a Marianas spinning disk imaging system with a Yokogawa CSU-W1 scanning unit on an inverted Zeiss Axio Observer Z1 microscope controlled by SlideBook 6 (Intelligent Imaging Innovations, Inc.). Images were acquired using a Photometrics Evolve, back-illuminated EMCCD camera (512 x 512 pixels) and a 100x (NA 1.4 oil, Plan-Apochromat, M27) objective.

The confocal microscope used was a laser scanning confocal microscope LSM880 (Zeiss) equipped with an Airyscan detector (Carl Zeiss) and a 40x oil (1.4) objective. The microscope was controlled using Zen Black (2.3) and the Airyscan was used in standard super-resolution mode.

The structured illumination microscope (SIM) used was DeltaVision OMX v4 (GE Healthcare Life Sciences) fitted with a 60x Plan-Apochromat objective lens, 1.42 NA (immersion oil RI of 1.516) used in SIM illumination mode (five phases x three rotations). Emitted light was collected on a front-illuminated pco.edge sCMOS (pixel size 6.5 mm, readout speed 95 MHz; PCO AG) controlled by SoftWorx.

### 2D Random Migration Assay

Cells were seeded at a density of 5 x 10^**3**^ in 1 ml media supplemented with 50 mM HEPES on plates coated with 10 μg/ml fibronectin and incubated for 2 hr at 37°C, 5% CO_2_. Live cell imaging was performed on a Nikon Eclipse Ti2-E widefield microscope with a heated CO_2_ chamber, Hamamatsu scientific CMOS Orca Flash 4 v4 and 10x Nikon CFI Plan Fluor objective. Random migration of cells was measured over 24 hr in a time-lapse movie with images taken every 10 min. Manual tracking of cells was performed using Fiji ImageJ MTrackJ plugin. The tracked data was analysed using Ibidi chemotaxis and migration tool to determine migration speeds, directionality and distance. Graphs of resulting data were produced using PlotsOfData (Postma and Goedhart, 2019). Randomization tests were performed using the online tool PlotsOfDifferences (https://huygens.science.uva.nl/PlotsOfDifferences/) (Postma and Goedhart, 2019).

### Quantitative RT-PCR

Total RNA extracted using the NucleoSpin RNA Kit (Macherey-Nagel) was reverse transcribed into cDNA using the high-capacity cDNA reverse transcription kit (Applied Biosystems) according to the manufacturer’s instructions. The RT-PCR reactions were performed using predesigned single tube TaqMan gene expression assays and were analysed with the 7900HT fast RT-PCR System (Applied Biosystems). Data were studied using RQ Manager Software (Applied Biosystems).

## Acknowledgements

We thank David Critchley for critical reading of the manuscript and Anthony Baines for his help with the bioinformatics of the *TLNRD1* gene. We also thank Soleil synchrotron beamline Proxima-1 staff for their help in crystallographic data collection. We thank J. Siivonen and P. Laasola for technical assistance and M. Saari for help with the microscopes. The Cell Imaging and Cytometry Core facility (Turku Bioscience, University of Turku, Åbo Akademi University and Biocenter Finland) is acknowledged for services, instrumentation, and expertise. B.T.G. was funded by BBSRC grants (BB/N007336/1 and BB/S007245/1) and B.T.G. and A.A were funded by HFSP grant (RGP00001/2016). G.J. was supported by grants awarded by the Academy of Finland, the Sigrid Juselius Foundation and Åbo Akademi University Research Foundation (CoE CellMech) and by Drug Discovery and Diagnostics strategic funding to Åbo Akademi University.

## Supplementary

**Figure S1.**
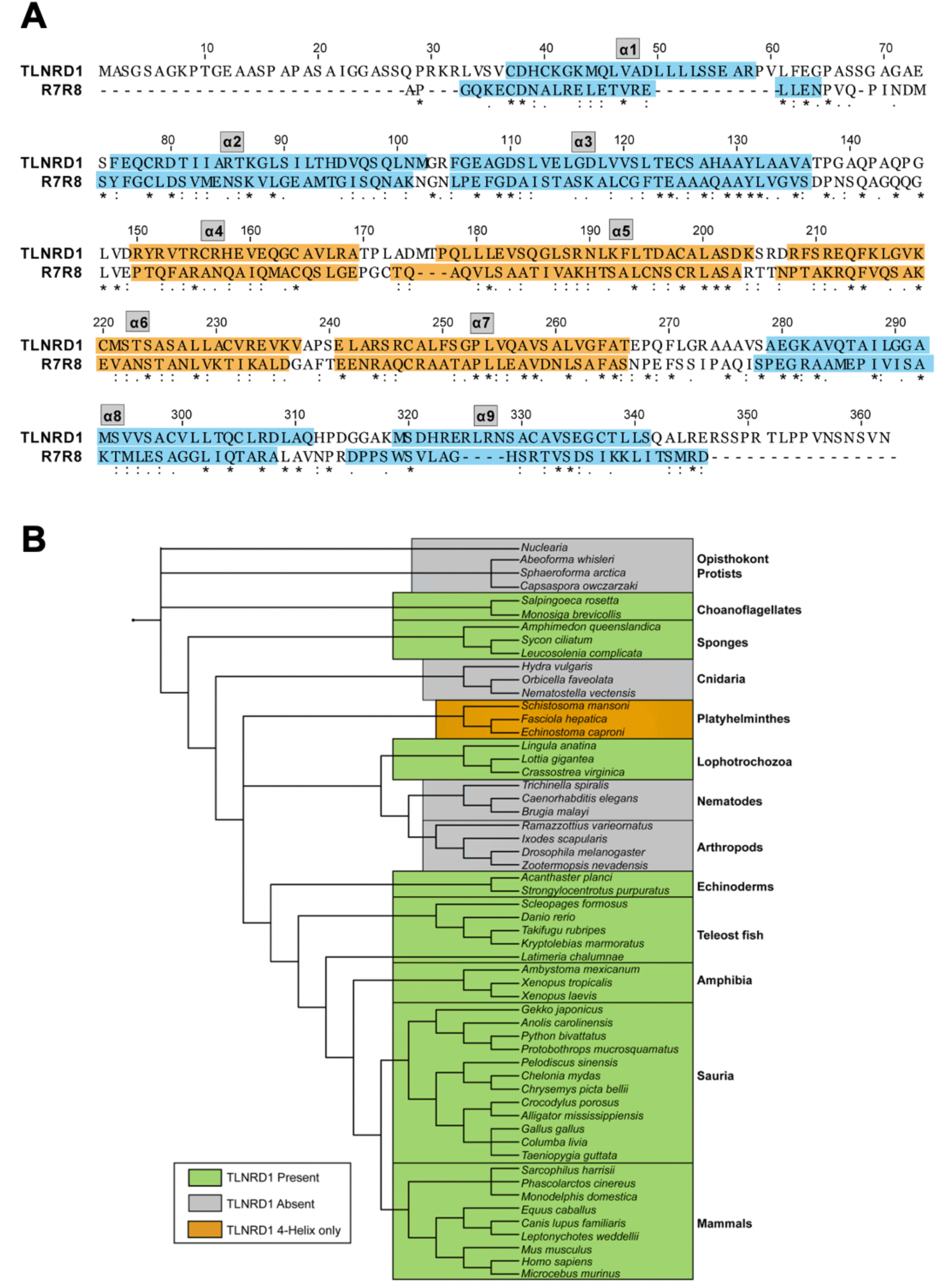
**(A)** Amino acid sequence alignment of human TLNRD1 (UniProt ID: Q9H1K6) and talin1 residues 1357-1653 encompassing R7R8 (UniProt ID: Q9Y490). Domain boundaries are highlighted with orange corresponding to the 4-helix domain and blue corresponding to the 5-helix domain. **(B)** Tree diagram representing TLNRD1 presence and absence over evolution. TLNRD1 first appears in choanoflagellates and sponges and is conserved throughout vertebrate evolution. TLNRD1 has been lost from Nematodes, Arthropods and Cnidaria.

**Figure S2.**
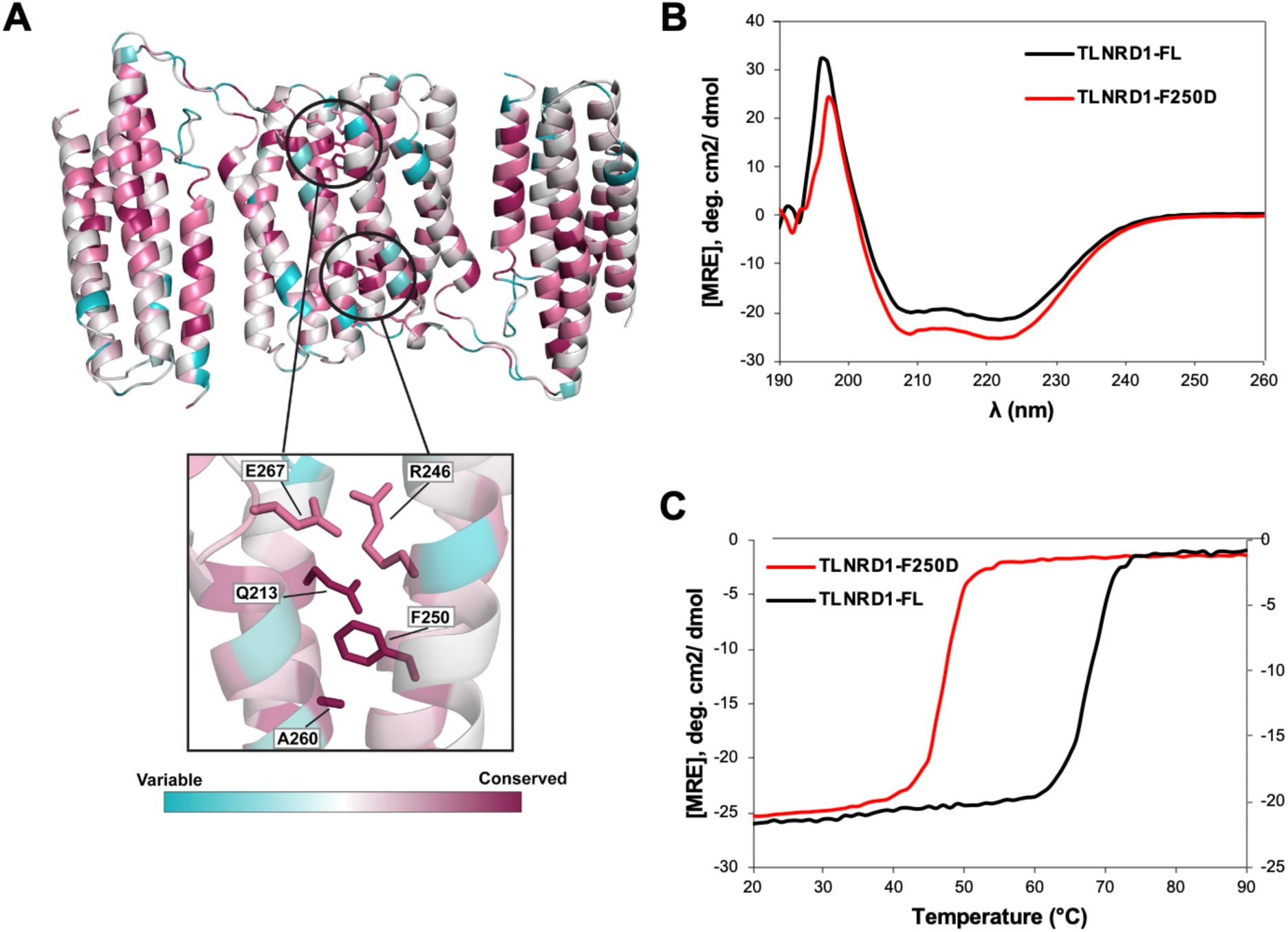
**(A)** TLNRD1-FL sequence conservation mapped onto the full-length TLNRD1 structure using ConSurf (Ashykenazy et al., 2016). Structure coloured according to extent of conservation with blue indicating variability and purple indicating highly conserved residues. Residues important for dimerisation (Q213, R246, F250, A260, E267) are highlighted. **(B)** Circular dichroism spectra of TLNRD1-FL (black) and TLNRD1-F250D (red) showing no change in overall secondary structure. **(C)** Circular dichroism thermostability analysis of TLNRD1-FL and TLNRD1-F250D showing reduction in stability with loss of dimerisation.

**Figure S3.**
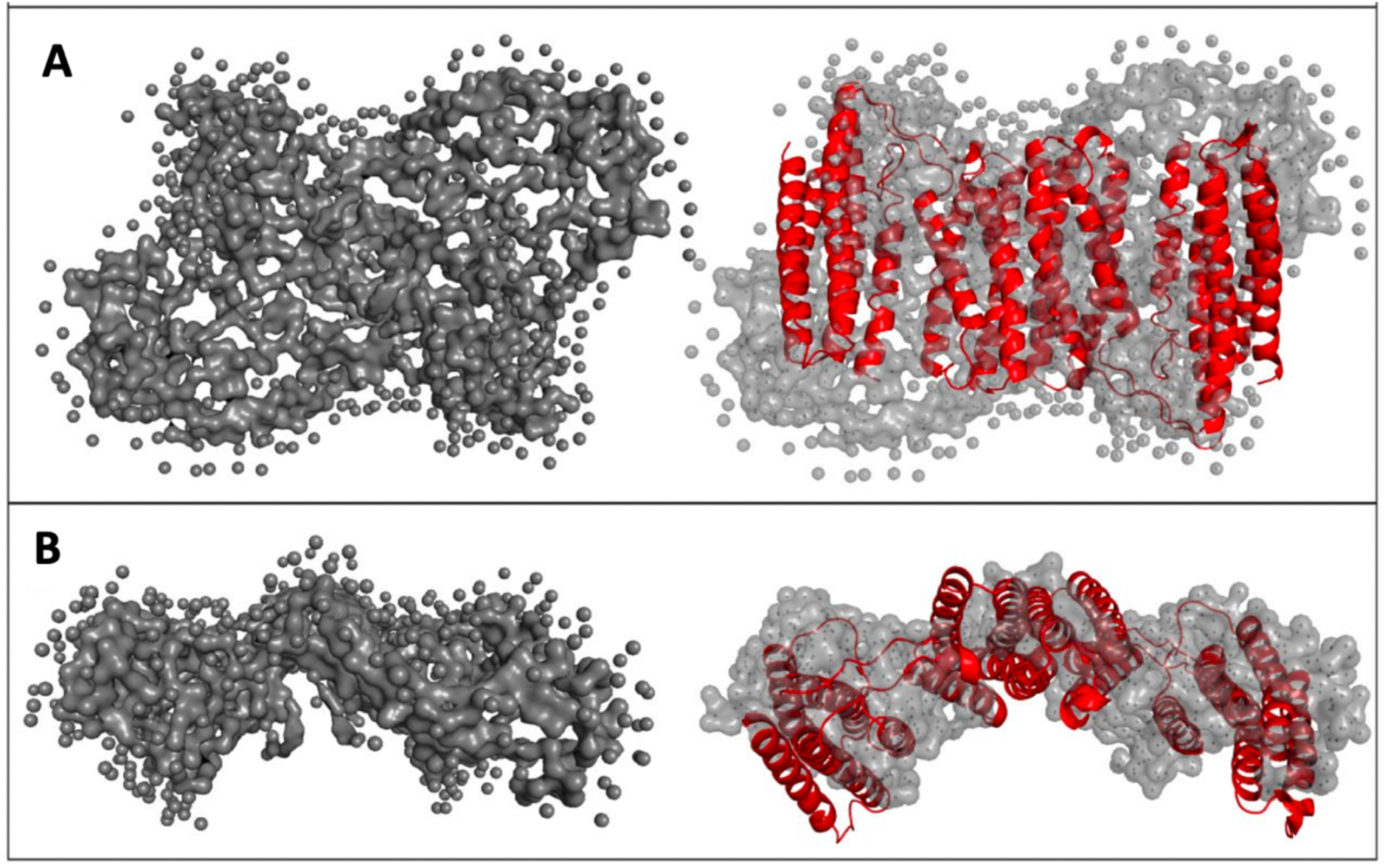
Small Angle X-Ray Scattering (SAXS) analysis of TLNRD1 shows compact domain arrangement in solution. **(A-B)** SAXS envelope reconstruction with GASBOR, showing **(A)** the best fit with the TLNRD1-FL crystal structure. **(B)** Top-down view of (A).

## Notes

### Competing Interest Statement

The authors have declared no competing interest.

